# Mapping the diffusional landscape of short NEAT1 in living cells

**DOI:** 10.64898/2026.05.13.724860

**Authors:** Sabrina Zappone, Eleonora Perego, Eli Slenders, Alberto Diaspro, Michele Oneto, Murat Sunbul, Giuseppe Vicidomini

## Abstract

The long non-coding RNA NEAT1 is a fundamental architect of nuclear condensates, specifically paraspeckles. While the scaffold-essential isoform NEAT1-2 has been extensively characterized, the function and dynamics of its shorter isoform, NEAT1-1, remain poorly understood. Investigating NEAT1-1 in live cells has been historically hindered by its genomic overlap with NEAT1-2. Traditional visualization study designs require either the genetic ablation of NEAT1-2, which disrupts paraspeckle integrity, or the use of bulky tandem tagging arrays, which can sterically hinder RNA folding and partitioning. Here, we implemented a non-invasive imaging strategy and performed diffusivity analysis of NEAT1-1 using the fluorescence light-up aptamer biRhoBAST. This small, high-affinity RNA tag enables high-contrast visualization of NEAT1-1 while preserving the structural integrity of both isoforms and their associated nuclear bodies. By combining imaging and fluorescence fluctuation spectroscopy, we provide characterization of NEAT1-1 within intact micro-and para-speckles. Our results reveal that NEAT1-1 is not purely sequestered within visible condensates; rather, a fraction exists in a distinct diffusive state within the nucleoplasm, likely as nanoscale complexes. These findings suggest that NEAT1-1 possesses a previously unrecognized regulatory role independent of the primary paraspeckle scaffold, offering new insights into the functional diversity of the lncRNA isoforms.

## 1. Introduction

The nucleus is organized by a diverse array of membrane-less organelles (MLOs), which are biomolecular condensates that concentrate specific RNAs and proteins without a lipid bilayer [1]. Many nuclear MLOs assemble around long non-coding RNAs (lncRNAs), which serve as structural backbone that recruit multiple RNA-binding proteins and nucleate higher-order assemblies [2]. A well-characterized example is the lncRNA NEAT1, which drives the formation of paraspeckles, nuclear bodies that regulate gene expression through the retention of edited transcripts [3]. The human *NEAT1* gene produces two isoforms from the same locus: the shorter NEAT1-1 (3.7 kb) and the longer NEAT1-2 (22.7 kb). NEAT1-2 serves as the essential architectural scaffold of paraspeckles, adopting a defined spatial organization that nucleates and maintains the entire structure [4]. In contrast, NEAT1-1 lacks the terminal domains required for strong interaction with essential paraspeckle proteins (PSPs) [5, 6] and behaves as a client molecule – a component that partitions into a liquid-like condensate without being required for its formation [7]. Despite being more abundantly expressed than NEAT1-2, NEAT1-1 has often been considered dispensable for normal cell homeostasis [8, 9]. However, emerging evidence points to distinct roles in cancer biology, where NEAT1-1 has been linked to both tumor-suppressive and oncogenic activities depending on cancer type and context [10–12].

Recent work has begun to address this gap by showing that NEAT1-1 localizes to a distinct class of nuclear bodies termed microspeckles. Microspeckles are significantly more numerous than canonical paraspeckles, persist in the absence of NEAT1-2, and concentrate at the peripheral zones surrounding nuclear speckles – the major hubs of pre-mRNA splicing and processing [5, 13]. Their consistent presence and defined positioning in the nucleoplasm argues against a non-functional role and points toward paraspeckle-independent functions. Despite these observations, the function of NEAT1-1 outside paraspeckles has remained entirely uncharacterized, and microspeckles have never been studied in living cells under physiologically intact conditions.

This gap is largely a consequence of a fundamental tagging or visualization problem. Because NEAT1-1 and NEAT1-2 share the same 5^′^ sequence, any probe targeting the 5^′^ region of NEAT1-1 inevitably co-labels NEAT1-2. Selective visualization of NEAT1-1 has therefore required the prior removal of NEAT1-2 expression, which simultaneously disrupts paraspeckles and eliminates the physiological nuclear context [5, 6]. RNA fluorescence in situ hybridization (FISH) suffers from the same limitation, as any probe designed against NEAT1-1 inevitably labels NEAT1-2 as well (fig. 1a). As a result, the behavior of NEAT1-1 across its distinct nuclear compartments – paraspeckles, microspeckles, and the surrounding nucleoplasm – has remained inaccessible in living cells.

**Figure 1.**
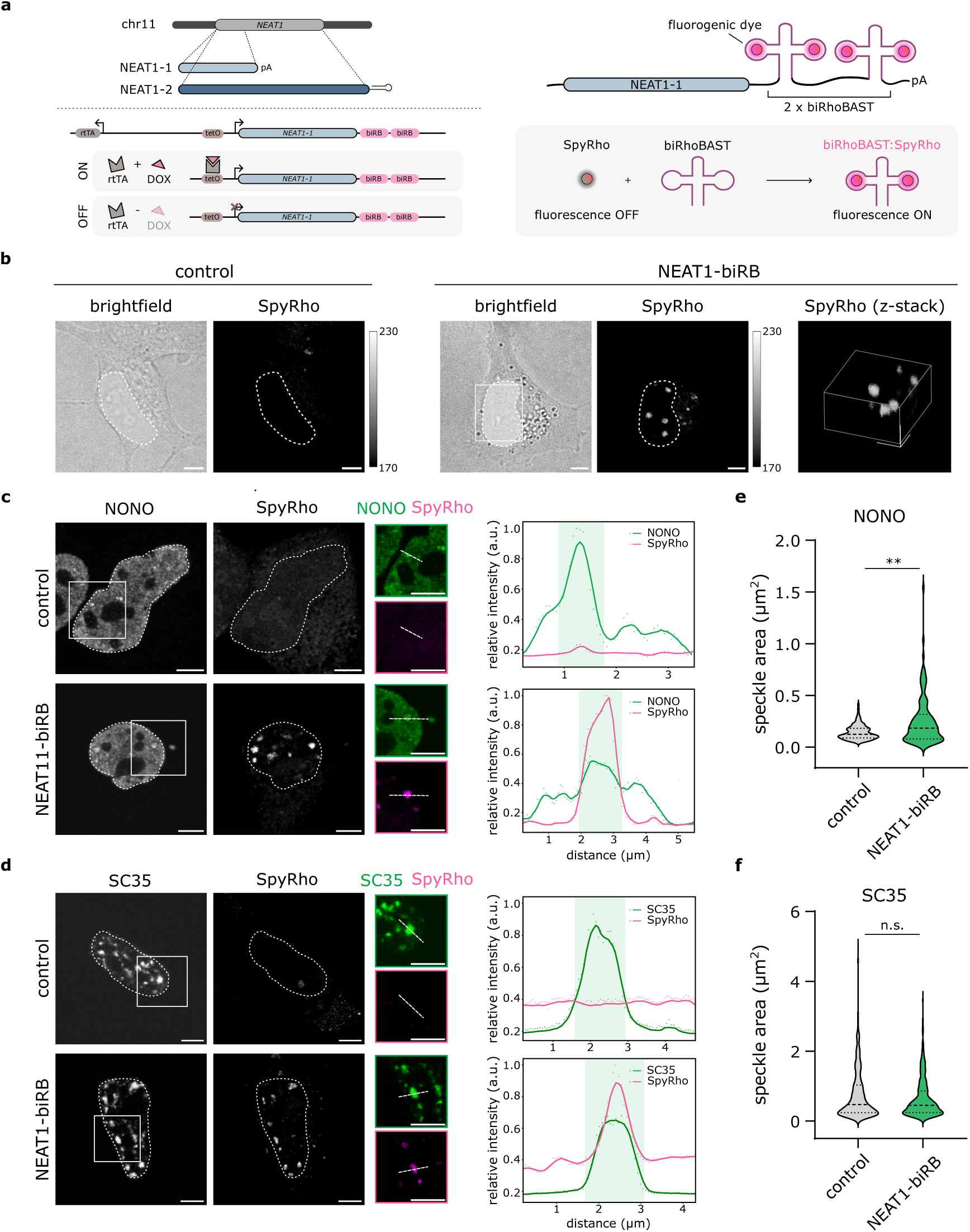
biRhoBAST-labeled NEAT1-1 localizes to speckles in live and fixed cells. **(a)** Schematic representation of the *NEAT1* gene, the tet-ON system used for NEAT1-biRB expression, and the biRhoBAST:SpyRho labeling strategy. **(b)** Representative brightfield and fluorescence images of SpyRho staining in live HeLa cells. Comparison between control and NEAT1-biRB-transfected cells demonstrates specific RNA labeling. Circles display the nuclear area. On the right, 3D reconstruction of z-stack (SpyRho signal) of the field-of-view indicated by the white square. Scale bar: 5 *µm*. **(c, d)** Colocalization of paraspeckle marker NONO (c) and nuclear speckle marker SC35 (d) with NEAT1-biRB (SpyRho). On the left, representative confocal images show untransfected control (top) and NEAT1-biRB transfected (bottom) cells labeled with SpyRho. Brightness and contrast were adjusted uniformly across the same channel for all images. On the right, plots display relative fluorescence intensity of line profiles across the indicated zoomed regions (white boxes). Scale bar: 5 *µm*. **(e, f)** Violin plots representing the distribution of paraspeckle (e) and nuclear speckles (f) areas in control and transfected (NEAT1-biRB) cells. The sample size (*n*) is: *n_ctrl_* = 108 and *n_NEAT_* _1_ = 102 for (e), *n_ctrl_* = 522 and *n_NEAT_* _1_ = 324 for (f). Horizontal lines indicate the median and 95% confidence intervals. Statistical significance was determined by a Mann-Whitney test: *p* = 0.005 (**) for (e) and *p* = 0.330 (n.s.) for (f).

Circumventing this limitation requires using genetically encoded tags that can be inserted at NEAT1-1–specific positions and read out directly in living cells. The most widely used system, MS2-MCP, implies the insertion of bulky tandem hairpin arrays into the RNA of interest and the co-expression of a fluorescent protein fused to an RNA-binding coat protein recognizing these hairpins. These systems can perturb the folding, localization, and function of the tagged RNA. Fluorescence light-up aptamers (FLAPs) offer a less invasive alternative: they are genetically encoded RNA aptamers that bind a cognate fluorogenic dye and activate its fluorescence, enabling live-cell RNA visualization without protein co-expression [14–16]. FLAPs such as Mango have already demonstrated their potential for labeling NEAT1-1 [17]. However, they generally suffer from limited brightness and residual fluorescence background from unbound dye. The bivalent aptamer biRhoBAST partially overcomes these limitations through improved binding affinity and effective brightness [18, 19]. Its RNA scaffold contains two binding sites that can engage either monovalent rhodamine ligands or dimeric variants in which two rhodamines are tethered together. With dimeric ligands, simultaneous engagement of both sites by a single molecule increases avidity and further enhances brightness and fluorogenicity [19, 20]. Among the available ligands, the spirocyclic rhodamine SpyRho combines high affinity with fast exchange kinetics, enabling the first super-resolved images of FLAP-labeled RNAs in live mammalian cells [20].

Although FLAPs allow for selective NEAT1-1 visualization, fluorescence imaging reports primarily on steady-state localization but not on the diffusional and interaction dynamics that govern MLO function. MLOs are dynamic environments whose functional state depends not only on molecular composition but also on how their components move, interact, and partition between microenvironments [21]. The diffusional landscape of an RNA – how freely it moves, how strongly it is confined, and whether it is transiently trapped in sub-resolution structures – is a key determinant of nuclear body function that imaging cannot capture. Accessing this information requires techniques that operate at nanoscale and report directly on local confinement architecture.

Fluorescence fluctuation spectroscopy (FFS) is a family of techniques that extract molecular parameters from fluorescence intensity fluctuations recorded in a diffraction-limited detection volume, such as the detection volume of a confocal microscope [22]. Within this family, fluorescence correlation spectroscopy (FCS) provides access to diffusion coefficients, interactions and molecular concentrations [23, 24]; spot-variation FCS (svFCS), which extends FCS measurements across systematically varied observation volumes, additionally yields the confinement mode, distinguishing free diffusion from transient nanodomain trapping and meshwork hop diffusion [25, 26]; and fluorescence lifetime analysis provides an orthogonal spectroscopic tool that can discriminate molecular nano-environment, conformation or binding of fluorophores on the basis of their excited-state decay [27]. Together, these modalities provide a multi-parameter readout of RNA mobility, confinement, and molecular identity at nanoscale [28–30]. We have previously shown that all these modalities can be acquired simultaneously from a single intensity measurement using photon-resolved microscopy based on a single-photon avalanche diode (SPAD) array detector [30], and applied this approach to decode molecular dynamics within multiple MLO systems [31–33]. Moreover, FFS is sensitive to nanomolar fluorophore concentrations [34], making it particularly suited to characterizing RNA dynamics in MLOs where expression levels and fluorophore brightness can be limiting. Finally, the fluorescence lifetime of the aptamer-bound dye is expected to differ from that of the unbound dye, offering a potential spectroscopic parameter that, when combined with FCS, can separate RNA-associated signal from background.

FFS has previously been applied to RNA aptamer systems, but only *in vitro* [35, 36] or with synthetic RNA-based tools [37]. Its application to a lncRNA like NEAT1-1 in living cells has not been reported. Here, we tagged NEAT1-1 with a minimal biRhoBAST cassette and combined this reporter with a multi-parameter SPAD array-based FFS platform to characterize NEAT1-1 mobility across distinct nuclear compartments – paraspeckles, microspeckles, and the surrounding nucleoplasm – dissecting its diffusional behavior in each. Our measurements reveal that NEAT1-1 encounters structurally distinct microenvironments as it diffuses through the nucleoplasm. This points to a model in which nucleoplasmic NEAT1-1 partitions into nanocondensates that may function as satellite assemblies, revealing an organizational dimension of NEAT1 biology that extends beyond its canonical paraspeckle role.

## 2. Results

### Validation of the NEAT1-biRhoBAST system via imaging

To enable live-cell imaging of NEAT1-1, we first engineered a doxycycline-inducible mammalian expression vector for the lncRNA isoform tagged with the fluorogenic aptamer biRhoBAST. Having multiple FLAP repeats can significantly enhance the signal-to-noise ratio, an essential requirement for low-abundance transcripts such as NEAT1-1 [18]. However, large tandem arrays risk perturbing the natural folding and functionality of the RNA. To mitigate these potential artifacts while ensuring sufficient detectable fluorescence, we constrained the tag to two biRhoBAST repeats placed at the 3’ terminus of the NEAT1-1 sequence (fig. 1a).

We selected HeLa cells as our primary model given their prominent paraspeckle formation and their established use for studying paraspeckle assembly [8]. To visualize NEAT1-1, we expressed a NEAT1-1 variant tagged with the aptamer (hereafter referred to as NEAT1-biRB) and stained cells with the biRhoBAST ligand SpyRho. EGFP (or nls-EGFP) was co-transfected as a transfection control. Following doxycycline induction, we observed the specific accumulation of NEAT1-biRB into distinct nuclear foci in transfected cells (fig. 1b). Although SpyRho is fluorogenic and should emit fluorescence only upon binding its aptamer, we assessed potential intracellular background by imaging non-transfected cells as a control. No significant fluorescence was detected in either control or non-induced (no doxycycline) cells (fig. 1b, S1a). To further exclude the possibility that observed speckle pattern was a SpyRho-specific artifact, we repeated the staining with an alternative biRhoBAST-compatible ligand, bivalent tetramethylrhodamine (TMR_2_) [19], and obtained equivalent localization (Fig. S1b,d).

Finally, to ensure that the system performance is cell-line independent, we further validated the NEAT1-1 imaging in HEK293T cells, observing consistent subcellular localization patterns across both models (fig. S1b,c).

As NEAT1-1 is expected to localize within two distinct types of nuclear bodies, we verified its recruitment to both paraspeckles and microspeckles. To assess paraspeckle localization, we performed immunofluorescence for the canonical marker NONO in transfected cells. Since the biRhoBAST system is compatible with both live and fixed-cell imaging [19], we compared the paraspeckle staining with SpyRho signal coming from NEAT1-biRB transcripts. Colocalization analysis between NONO and SpyRho signals, supported by line profile plots, confirmed that overexpressed NEAT1-biRB was successfully recruited into endogenous paraspeckles (fig. 1c). While the proteomic composition of paraspeckles is well-characterized, that of microspeckles, where NEAT1-1 isoform is known to localize, remains poorly defined. These structures are currently identified by their spatial proximity to nuclear speckles, marked by SC35. Although SC35 and NEAT1-1 reside in distinct MLOs, the separation between microspeckles and nuclear speckles falls below confocal spatial resolution and produces near-complete signal overlap [5]. We therefore verified the recruitment of NEAT1-biRB to these nuclear bodies by analyzing the colocalization between SpyRho and SC35 signals (fig. 1d), confirming that our engineered NEAT1-1 correctly partitions into both NONO-positive paraspeckles and SC35-proximal microspeckles.

The specificity of NEAT1-1 spatial localization was further corroborated by analyzing the morphological response of nuclear bodies to its overexpression. Nuclear speckle (SC35-positive) size has been shown to scale with the abundance of their integrated RNA components [38]. Because NEAT1-1 is not an endogenous component of nuclear speckles, its overexpression and the potential increase in condensate recruitment should not alter SC35 speckle dimensions, even if peripheral microspeckles accumulate nearby. Consistent with this prediction, we observed no significative change in nuclear speckle area upon NEAT1-1 overexpression compared to untransfected control cells (fig. 1f). These speckles retained their characteristic morphology, with diameters ranging from 0.5 *µm* to few micrometers and a granular, interconnected structure [13, 38]. A complementary question is whether NEAT1-1 affects paraspeckles, the bodies in which it does partition. Previous studies indicated that NEAT1-1 cannot independently regulate the size or number of paraspeckles in the absence of NEAT1-2 [5]. Our system enable to perturb NEAT1-1 levels while leaving NEAT1-2 intact, thus allowing us to isolate the effect of NEAT1-1 on pre-existing paraspeckles. In control cells, paraspeckle number and area fell within the range reported for HeLa cells (10–30 paraspeckles per nucleus) [8, 39]. Upon NEAT1-1 overexpression, both paraspeckle features increased significantly, despite the relatively weak interaction of NEAT1-1 with essential PSPs [5] (fig. S3c, 1e). Beyond the functional implications of this volume change, these morphological measurements provide together a two-sided validation of our system: NEAT1-biRB selectively enlarges only the bodies in which it accumulates (paraspeckles) while leaving unrelated MLOs (nuclear speckles) unchanged. Together with the colocalization with paraspeckle markers, this selectivity confirms that NEAT1-biRB reports on NEAT1-1 subcellular localization.

We recognized that overexpression of NEAT1-1 and the subsequent increase in condensate RNA content could, under certain circumstances, alter the physical state of MLOs [40]. To confirm that NEAT1-1 nuclear bodies retain their liquid-like properties, we performed a 1,6-hexanediol (1,6-HD) assay. As an aliphatic alcohol, 1,6-HD is a widely utilized chemical probe for investigating the material properties and structural integrity of biomolecular condensates. It acts by disrupting the weak hydrophobic interactions that stabilize these liquid-like assemblies, leading to reversible dissolution of MLOs while leaving solid-like aggregates intact or less sensitive to dissolution [6, 41, 42]. Following incubation of both control and transfected cells with 1,6-HD, we observed the near complete dissolution of paraspeckles (marked by NONO) and any visible speckle (fig. S3). These results confirm that our experimental conditions do not shift the material state of NEAT1-1 assemblies toward solid-like assemblies, demonstrating that NEAT1-enriched bodies maintain the characteristic liquid-like behavior of physiological MLOs.

### Validation of the NEAT1-biRhoBAST system via spot-variation FCS

Having confirmed the spatial recruitment of NEAT1-biRB to nuclear sub-compartments via imaging, we next characterized its mobility. To this end, we employed a framework based on FCS [28]. Specifically, we recorded the intensity fluctuations of SpyRho in distinct nuclear regions (nucleoplasm and NEAT1-1 foci) in both control and NEAT1-biRB-expressing cells. In FCS, the temporal autocorrelation of these intensity signals allows for the extraction of parameters such as the diffusion coefficient of a fluorescent reporter. The diffusion coefficient is derived from the characteristic diffusion time (*τ_D_*), defined as the average residence time of a fluorophore within the confocal detection volume [22]. *τ_D_* is determined by fitting the autocorrelation function (ACF) to a predefined mathematical model. However, choosing a specific model often necessitates prior knowledge of the local diffusivity landscape or the number of diffusing components, which can introduce significant bias into the analysis, especially for a highly heterogeneous system such as the nuclear environment. To circumvent this bias, we first applied the Maximum Entropy Method (MEM) analysis to the measured ACFs. MEM is a powerful numerical approach that fits the ACF with a free diffusion model assuming the presence of hundreds of components, yielding a histogram representing the distribution of diffusion times *τ_D_* in the sample [43]. Based on the number of discrete peaks identified in the MEM histogram, we subsequently applied a one- or two-component free diffusion model to fit the ACFs (fig. 2a). This allowed us to retrieve the apparent diffusion coefficient (*D_app_*) for SpyRho across different nuclear environments.

**Figure 2.**
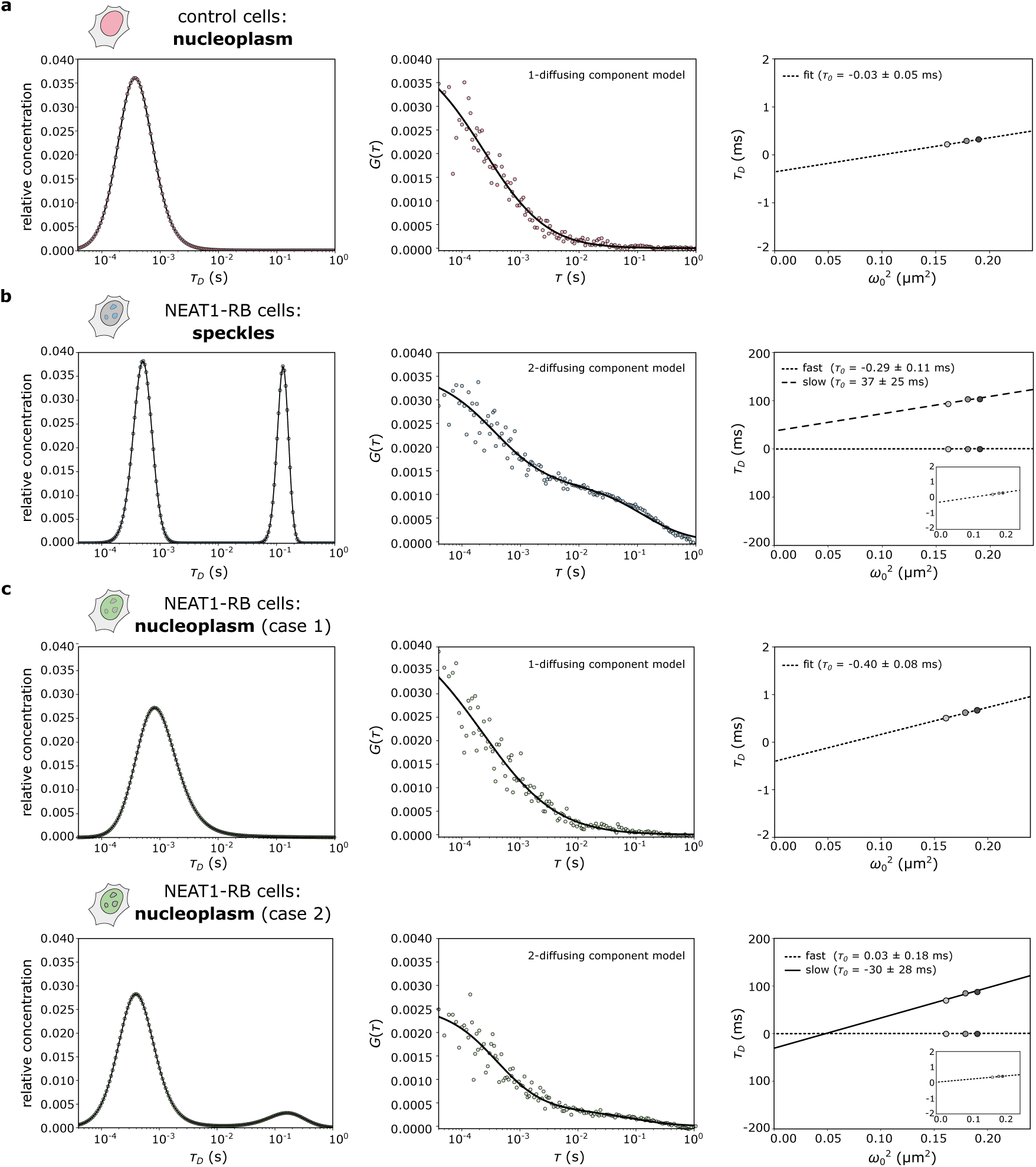
biRhoBAST:SpyRho reveals distinct NEAT1 dynamics across cellular compartments. Representative FCS measurements of SpyRho in control (a) and in transfected cells recorded inside (b) or outside (c) nuclear speckles. Each row shows, from left to right, the MEM analysis used to identify the diffusing species, the FCS autocorrelation curve *G*(*τ*) with its corresponding fit, and the diffusion law 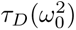 from which the intercept *τ*_0_ is determined. In (b) and (c), insets show the diffusion law of the fast component on a rescaled y-axis. Fitting models for 1-or 2-diffusing components were applied based on the number of peaks identified in the MEM distribution. For nucleoplasmic measurements (c), examples of both 1-and 2-component scenarios are shown to illustrate the heterogeneity observed in this compartment.

While *D_app_*quantifies the diffusion, it does not inherently provide an overview of the local micro-environment (i.e., diffusion mode) influencing molecular movement. Traditionally, diffusion modes are determined by performing sequential FCS measurements at varying observation volumes (the so-called svFCS) [25]. The resulting plot of *τ_D_ versus* observation area (*ω*^2^) yields a linear relation called diffusion law *τ_D_*(*ω*^2^) (fig. S2b-c). The y-intercept of this linear fit (*τ*_0_) serves as a parameter to distinguish between different diffusion modalities (Brownian, confined, and hop diffusion). Specifically, a null intercept (*τ*_0_ = 0) indicates Brownian motion, while a positive intercept (*τ*_0_ *>* 0) indicates confined motion, and a negative intercept (*τ*_0_ *<* 0) is characteristic of meshwork or hop diffusion through a partitioned environment [25, 26] (fig. S2b). In dynamic systems such as living cells, the local molecular environment is temporally heterogeneous and can evolve between successive acquisitions. As a result, each data point in the diffusion law may sample a different instantaneous state (e.g., nanocondensates assemble and disassemble on timescales of seconds [44]). Our platform employs a single-photon-avalanche diode (SPAD) array detector which mimics distinct observation volumes simultaneously, ensuring that all points of the diffusion law are extracted from the same snapshot (fig. S2b). This approach allows us to determine the diffusion mode in a single measurement, thereby eliminating temporal non-stationarity and minimizing phototoxicity associated with repeated laser exposure on the same cellular spot [28].

By applying this analytical workflow to NEAT1-biRB:SpyRho system, we delineated distinct diffusion scenarios based on the local nuclear environment. In control cells, measurements were taken in random spots of the nucleoplasm (fig. 2a). Given the fluorogenic nature of SpyRho and the negligible background fluorescence detected in the nuclei of control cells, the presence of an ACF in this sample requires clarification. Certain cellular components may weakly and non-specifically activate SpyRho fluorescence, though with substantially lower efficiency than the biRhoBAST aptamer. Because FCS is sensitive to picomolar fluorophore concentrations, even a small fraction of fluorescent unbound dye is sufficient to generate a measurable ACF. The single diffusing component detected in control cells can therefore be attributed to this unbound SpyRho fraction. This interpretation is further proven by its diffusion modality; the diffusion law reveals a Brownian motion for this SpyRho population (fig. 2a, 3c). Due to its small hydrodynamic radius, a synthetic rhodamine such as SpyRho is indeed expected to diffuse into the nuclear environment without being significantly hindered by macromolecular crowding at the scales probed [45].

We performed the same FCS-based analysis on NEAT1-biRB-expressing cells, comparing regions both within and outside visible speckles. MEM analysis detected two diffusing components within NEAT1-1 speckles, whereas a mixed population was observed in the nucleoplasm (fig. 2b-c). Specifically, 65% of the nucleoplasmic measurements displayed a single diffusing component, while the remain exhibited two (fig. 2c). Importantly, *D_app_*values of the fast-diffusing component were comparable to those measured for unbound SpyRho in control cells (fig. 3b). In all tested conditions, this component exhibited Brownian motion as its diffusion mode (fig. 3c). To further validate our hypothesis that the fast-diffusing component represents unbound SpyRho, we removed the biRhoBAST repeats from NEAT1-biRB construct and induced NEAT1-1 overexpression alone. As expected, we detected only a single diffusing component (fig. S4b). This component exhibited a diffusion coefficient (*D_app_*(*control*) = 170±15 *µm*^2^*/s*; *D_app_*(*neat*1) = 148 ± 13 *µm*^2^*/s*) and diffusion modality (*τ*_0_(*control*) = −0.17 ± 0.08 *ms*; *τ*_0_(*neat*1) = −0.04 ± 0.09 *ms*) statistically indistinguishable from those of unbound SpyRho in control cells (fig. S4c).

**Figure 3.**
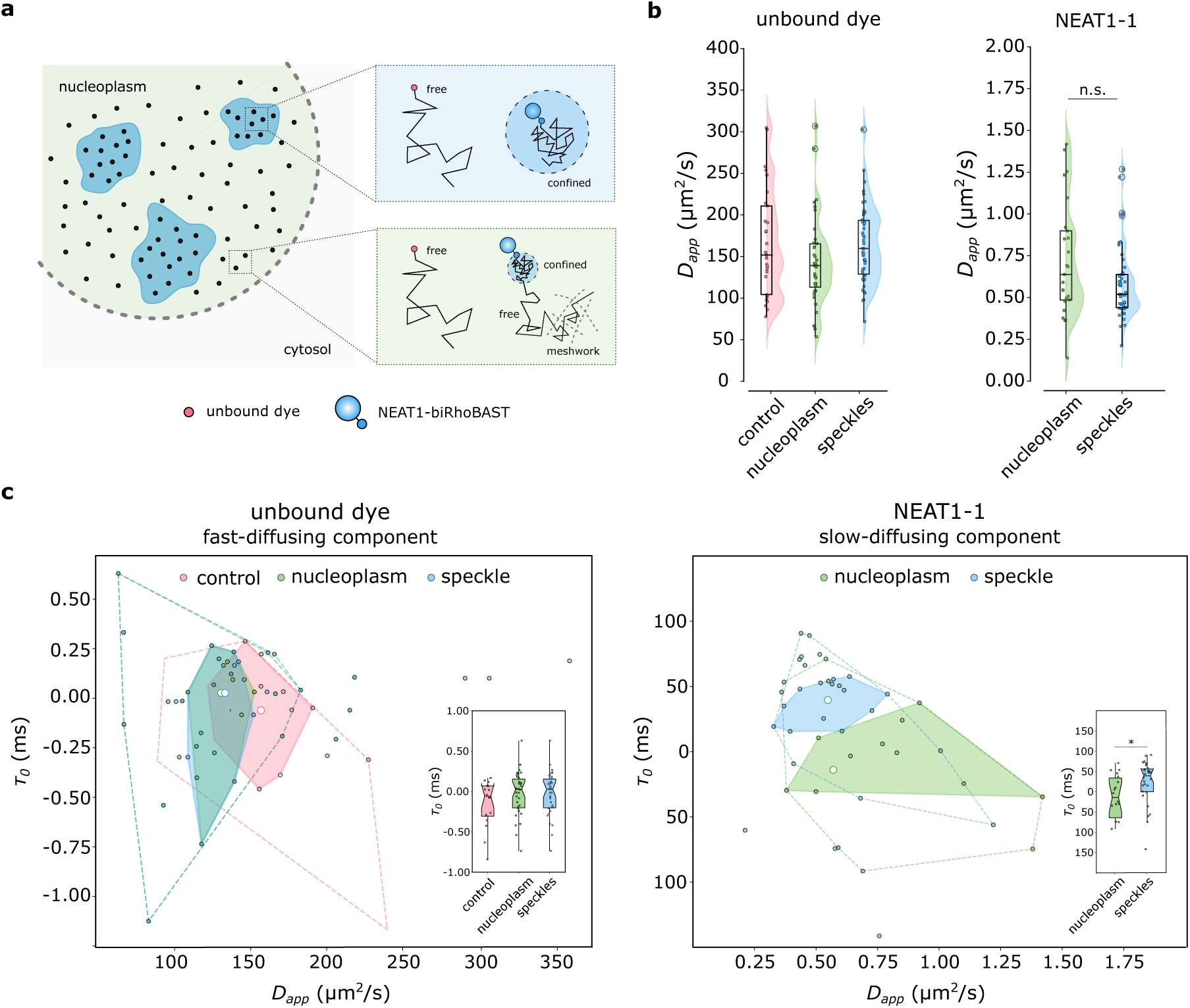
Comprehensive biRhoBAST:SpyRho measurements confirm distinct NEAT1-1 dynamics in nuclear subcompartments. **(a)** Graphical representation of the diffusional landscape of NEAT1-biRB within and outside speckles. **(b)** Distribution of the apparent diffusion coefficient *D_app_*for fast-diffusing and slow-diffusing components. Box plots and overlaid split violin plots compare control cells with the nucleoplasm and speckles of transfected cells. **(c)** Bag plots showing the correlation between *D_app_* and *τ*_0_ for unbound dye (left) and NEAT1-1 (right). Insets display notched plots of *τ*_0_ for each condition. In (c), statistical significance was determined by a Mann-Whitney test: *p* = 0.09 (n.s.) for *D_app_* and *p* = 0.02 (*) for *τ*_0_.

As an additional check, we asked whether the diffusion coefficient of the fast component was physically consistent with that of a free rhodamine dye in the nuclear environment. The nucleoplasmic viscosity (*η*) of HeLa cells is estimated to be 3-4 times larger than that of water at 37 ^◦^*C* [46]. Assuming *η*(*nucleus*) = 1.40 ± 0.27 *cP* [46] and a measured *D_app_*(*control*) = 170 ± 15 *µm*^2^*/s*, the calculated hydrodynamic radius is *r_H_*= 0.76 ± 0.19 *nm* (eq. 3). While a precise *r_H_* for SpyRho has not been reported, this value aligns closely with those of structurally similar rhodamines (e.g., *r_H_*= 0.77 *nm* for Rhodamine 110 [47]), supporting our identification of the fast-diffusing component as unbound dye.

Together, these observations confirm that the fast-diffusing component represents free, unbound dye. Since SpyRho exhibits fast ligand exchange kinetics and does not bind covalently to biRhoBAST, no washing steps were performed after incubation [20]. This likely results in a persistent pool of excess dye. In practice, our analysis effectively resolves this component in all conditions, ensuring that the presence of unbound SpyRho does not bias our subsequent characterization of NEAT1-1 mobility.

### Spot-variation FCS reveals NEAT1-1 dynamics in living cells

The slow-diffusing component was detected both within and outside visible speckles (fig. 2b-c). As this component is unique to NEAT1-biRB-expressing cells, it most likely represents SpyRho molecules bound to the biRhoBAST aptamer, thereby directly reporting on NEAT1-1 mobility. Its diffusion coefficient is more than two orders of magnitude lower than that of fast-diffusing component (fig. 3b). Because *D_app_* is sensitive to both molecular dimensions and local viscosity (eq. 3), this observed decrease likely reflects the transition of the dye from a small-molecule state to a bulky RNA-dye complex, along with the effective viscosity of the crowded nucleoplasm [48].

Inside visible speckles, NEAT1-1 exhibits strongly confined diffusion with a consistently positive *τ*_0_. This is indicative of diffusion restricted by microdomains, a behavior characteristic of molecules moving freely within local compartments but encountering energy barriers at their boundaries [25]. This is typical of dense condensates, where RNA and proteins create a network of transient binding sites that confine molecules below the diffraction limit [33, 49, 50]. Inside the nucleoplasm, the slow component could, in principle, represents NEAT1-biRB assemblies not detectable via confocal raster scanned imaging due to low RNA concentration or suboptimal brightness. However, the nucleoplasmic slow component displays a more heterogeneous landscape of diffusion modes, with a significantly shorter *τ*_0_ than speckles, suggesting that these assemblies represent a structurally distinct diffusional state (fig. 3c).

Despite this difference in confinement, both the speckle-associated and nucleoplasmic populations exhibit nearly identical *D_app_* values (fig. 3b). This suggests that the local microviscosity and the density of the immediate molecular environment are consistent across both structures, regardless of whether the NEAT1-1 molecules have assembled into larger, visible structures. In contrast, the lower *τ*_0_ in the nucleoplasm indicates that these sub-diffraction nanoclusters lack the rigid structural scaffolding or dense physical barriers characteristic of mature speckles, allowing for a more expanded spatial sampling of the local environment. This reflects structural or interaction differences rather than viscosity differences.

### Fluorescence lifetime discriminates between bound and unbound SpyRho

Beyond mobility studies, our photon-resolved microscope enables the simultaneous investigation of the photophysical properties of fluorophores diffusing within the confocal volume through fluorescence lifetime quantification. As a sensitive reporter of the local molecular environment, fluorescence lifetime provides a robust readout of distinct fluorophore photophysics states [27, 51]. Specifically, fluorophores used in FLAPs systems typically exhibit a significant longer fluorescence lifetime upon binding [52, 53]. This occurs because the RNA aptamer effectively shields the bound fluorophores from the solvent, suppressing non-radiative decay pathways and enhancing quantum yield (thus, resulting in longer lifetimes). For instance, FLAPs such as Riboglow [53], Mango, and Spinach [52] have been shown to extend the lifetimes of their respective ligands by few nanoseconds upon transitioning from the unbound to the bound state. In the case of SpyRho, the free dye exists predominantly in a non-fluorescent spirocyclic form whose photophysical properties differ fundamentally from the aptamer-bound open form [20], providing intrinsic spectroscopic contrast exploitable by fluorescence lifetime analysis.

We first measured the fluroscence lifetime of SpyRho within both control cells and NEAT1-biRB-expressing cells (speckles). Photon arrival time histograms were fitted with a monoexponential decay model to retrieve the fitted fluorescence lifetime (*τ_L_*): *τ_L_* = 2.68 ± 0.01 *ns* for control cells and *τ_L_* = 2.78 ± 0.01 *ns* for NEAT1-biRB speckles (fig. S5a, 4a). The modest shift (∼ 0.1 *ns*) between conditions suggested that, if aptamer-bound SpyRho carries a distinct lifetime, its contribution is largely masked by the dominant unbound population. In fact, in other FLAP–dye systems, binding induces lifetime differences of 3-4 ns between bound and unbound states, but the bound fraction may represent only a small proportion of the total fluorescent pool.

Given the characteristics of the biRhoBAST:SpyRho system, we hypothesized that SpyRho follows the same lifetime trends observed in other FLAP systems upon binding. This would account for the longer lifetime within speckle environment, where the presence of bound SpyRho is expected. To test whether a spectrally distinct bound population underlies this small average shift, we employed fluorescence lifetime fluctuation spectroscopy (FLFS), a combination of FCS and fluorescence lifetime measurement. While FLFS is often used to suppress detector artifacts (e.g., afterpulsing) [29], it is equally powerful for resolving coexisting lifetime populations in the same sample.

Instead of calculating a standard ACF on the raw photon counts, this analysis generates ACFs from lifetime-weighted photon traces. These weighted functions are obtained by assigning mathematical filters based on the distinct decay patterns of the species [29, 54]: one filter for unbound SpyRho (*τ_L_*(*short*)) and another for bound SpyRho (*τ_L_*(*long*)) (fig. S5). When a sample contains both lifetime populations, the analysis yields two distinct, species-specific ACFs. We systematically explored lifetime filtering windows centered on the two components and evaluated the quality of the resulting ACFs. Only a separation of *τ_L_*(*short*) = 2 *ns* and *τ_L_*(*long*) = 4 *ns* yielded well-defined, physically consistent ACFs for both populations (fig. 4b). Filtering with *τ_L_*(*short*) set to the fitted control *τ_L_*(2.68 ± 0.01 *ns*) or with narrower separations failed to resolve distinct diffusive species. The requirement for *τ_L_*(*short*) = 2 *ns* rather than the fitted value of 2.68 ± 0.01 *ns* likely reflects two factors. First, the fitted *τ_L_*is a single effective lifetime obtained under the assumption of mono-exponential decay. Unbound SpyRho in living cells likely samples a distribution of environments, so its true emission decay is unlikely to be strictly mono-exponential [55, 56]. In FLCS, what matters is the full decay pattern rather than a single lifetime value [57]. Because we define each filter using a single *τ_L_* value rather than a full multi-exponential pattern, filter performance depends on how well a mono-exponential decay approximates the actual species [54]. The fitted *τ_L_*may therefore be a suboptimal filter input even when it accurately summarizes the average decay. Second, effective lifetime filtering requires sufficient separation between the two components to discriminate photon populations: a small difference between *τ_L_*components degrades filter performance [54].

**Figure 4.**
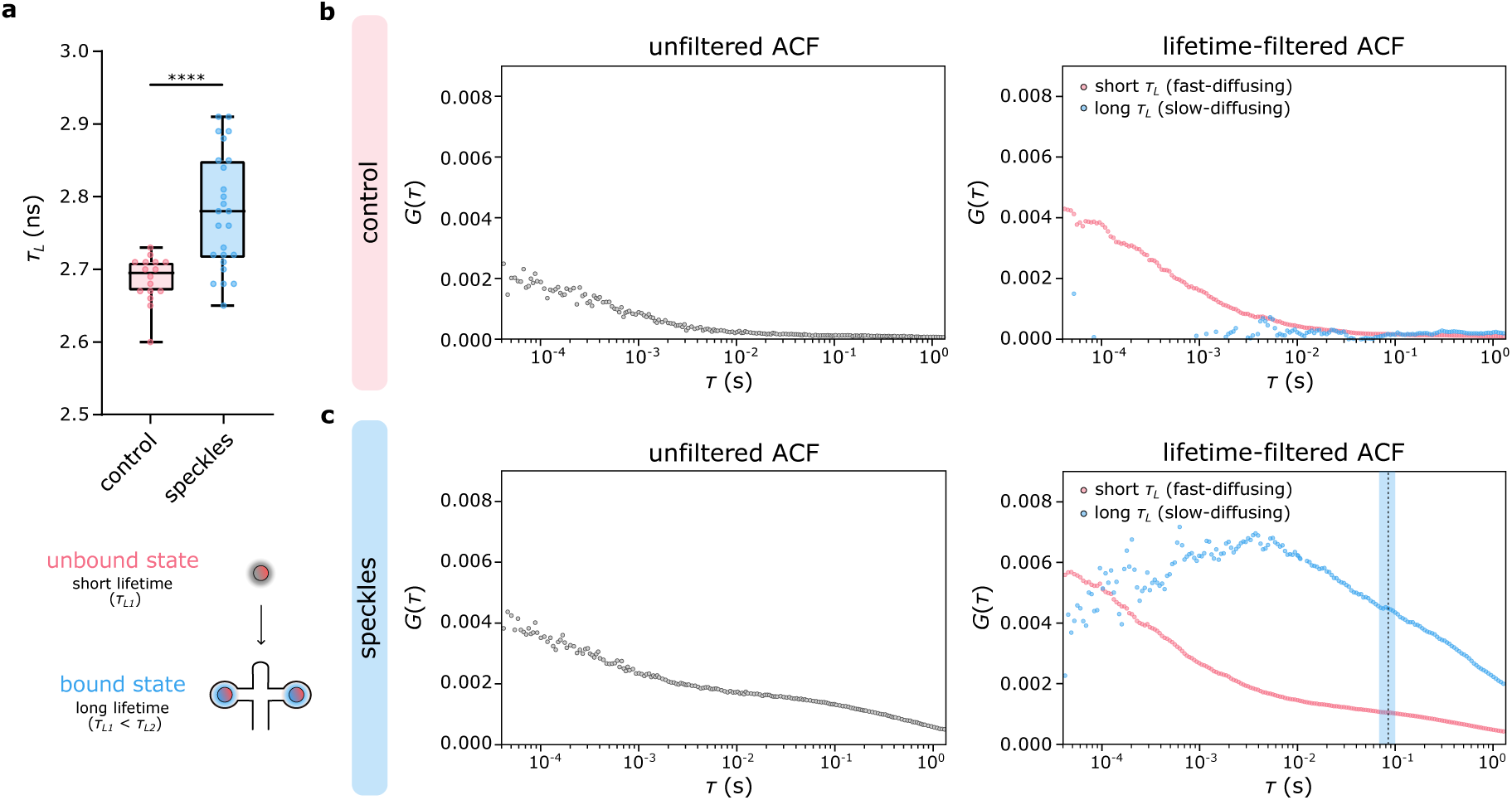
Bound and unbound SpyRho exhibit a different fluorescent lifetime. **(a)** Box plot of fluorescence lifetime values (*τ_L_*) extracted from FLFS measurements in control or NEAT1-biRB-expressing cells. The sample size is *N* = 16 for control and *N* = 25 for speckles. Statistical significance was determined by a T-test with Welch’s correction: *p <* 0.0001. **(b,c)** Autocorrelation functions *G*(*τ*) before (left) and after (right) lifetime-based filtering in control (b) and transfected (c) cells. Correlations corresponding to a longer *τ_L_*are in blue, the ones corresponding to a shorter *τ_L_* are in pink. Averaging of *n* = 10 and *n* = 12 measurements for control and transfected cells, respectively. For lifetime-filtered ACF in speckles, a vertical line shows *mean* ± *IQR* of *τ_D_* values presented in fig.2b for slow-diffusing component (speckles).

As expected, measurements in control cells yielded only one valid ACF (associated with *τ_L_*(*short*)), while no correlation was identified with the *τ_L_*(*long*) filter (fig. 4b). In contrast, measurements within speckles produced two distinct ACFs (fig. 4c). Although FLFS is intrinsically prone to noise due to a lower effective number of photons, we mitigated this by pooling multiple measurements. The resulting correlations for the NEAT1-biRB speckles clearly show two components. While the noise level makes fitting for precise *D_app_*values challenging, the ACF associated with *τ_L_*(short) is common to both conditions and represents the fast-diffusing, unbound dye. This aligns with our previous MEM-based characterization, which identified the unbound dye as the common diffusing component across all conditions. Furthermore, the speckle-specific ACF associated with *τ_L_*(long) exhibits a significantly higher diffusion time *τ_D_* (or lower *D_app_*). The absence of correlation at shorter timescales (i.e., *τ <* 1 *ms*) supports the conclusion that this component represents SpyRho bound to the bulky NEAT1-biRB complex (*τ_D_* = 84.7 ± 4.3 *ms*, fig. 2b).

## 3. Discussion

In this study, we applied a non-invasive fluorescent labeling approach to NEAT1-1 in living cells under endogenous conditions while preserving the structural scaffold NEAT1-2, enabling both imaging and nanoscale quantitative analysis of its mobility. By tagging NEAT1-1 with a minimal number of biRhoBAST repeats at its 3^′^ region, we designed a labeling strategy compatible with future endogenous tagging via gene editing, which cannot occur at the 5^′^ region and likely requires a small tag to preserve NEAT1-2 transcription. Validation through imaging and dynamic analyses confirmed that NEAT1-1 localization and function are not impaired by this tag, establishing a robust foundation for future studies of endogenous NEAT1-1-based biology.

In the context of NEAT1-1 overexpression, we observed that NEAT1-1 partitions into both the canonical paraspeckle core – likely by integrating into the pre-existing architectural framework provided by NEAT1-2 – and the more numerous, peripheral microspeckles. While NEAT1-2 serves as the essential architectural scaffold of paraspeckles, NEAT1-1 likely functions as a “client” molecule [7, 58], consistent with its lack of the terminal domains required to interact strongly with essential PSPs [5]. Accordingly, the observed increase in paraspeckle area upon NEAT1-1 overexpression most likely reflects a volume expansion of the paraspeckle “shell” driven by the accumulation of NEAT1-1 and its associated proteins, rather than *de novo* reorganization of the underlying NEAT1-2 “skeleton” [59]. Alongside this increase in paraspeckle area, we also observed higher number of paraspeckles per cell upon NEAT1-1 overexpression. It has been shown that paraspeckles can bud or split into clusters of multiple smaller structures around the NEAT1-2 transcription site once they reach a certain size [60]. Since budding alone cannot account for the simultaneous increase in both paraspeckle number and size [60], it is plausible that elevated NEAT1-1 levels also promote stochastic recruitment of paraspeckle-associated factors along newly transcribed NEAT1-2, facilitating biogenesis of additional paraspeckles. Collectively, these findings suggest that the marginal role previously attributed to NEAT1-1 in paraspeckle biology should be reconsidered.

To investigate the diffusional behavior of NEAT1-1 at the nanoscale, we combined FLAPs with FFS. We kept the number of biRhoBAST repeats minimal (only two) to limit any potential impact of the aptamer tag on the RNA of interest, a particularly important constraint for RNAs acting as scaffolds for MLO assembly. While two repeats represent one of the smallest FLAP tags currently available, they yield insufficient brightness for single-molecule imaging, precluding the study of RNA in small foci or in the dilute phase. Among the available FLAPs, biRhoBAST has shown a remarkable ability to image single RNA molecules by super-resolution microscopy, but single-mRNA tracking required at least eight repeats [19]. Single-particle tracking approaches offer high spatial resolution and can report on RNA mobility, but similarly depend on bright RNA complexes, posing a significant technical barrier. FCS and its variants, by contrast, are sensitive to picomolar fluorophore concentrations and therefore tolerate both low-brightness labels and low-abundance RNAs. We leveraged this sensitivity to characterize NEAT1-1 mobility at length scales inaccessible to imaging or conventional single-molecule techniques.

The high sensitivity of FFS, however, comes at a cost. The rhodamine ligand of biRhoBAST is not fully dark in its unbound state due to potential interactions with cellular components. While this fluorescence signal is negligible for imaging, free dye diffusing through the observation volume generates a fast-diffusing component in the svFCS correlation that contaminates the RNA signal and can bias the analysis of RNA diffusion. Here, we extended FFS to a FLAP-labeled endogenous lncRNA in living cells while accounting for the fluorescence background arising from unbound dye. Aptamer-bound dye often exhibits a different fluorescence lifetime than the free form. Since our platform allows svFCS and fluorescence lifetime detection in a single measurement, we developed an analysis pipeline that discriminates biRhoBAST-bound from unbound diffusion.

NEAT1-1 exists in two distinct diffusional states that share a common internal density but differ in their architectural constraints. Our data reveal that NEAT1-1 exhibits confined diffusion within speckles. At least for paraspeckles, this is consistent with transient confinement by the PSP network organized around the NEAT1-2 scaffold [61]. Outside speckles, NEAT1-1 shows a spatially heterogeneous, mixed distribution of *τ*_0_ values — ranging from positive to negative — while the diffusion coefficient *D_app_* remains identical to that measured inside MLOs. The uniform *D_app_* values imply that the local microviscosity experienced by NEAT1-1 is homogeneous throughout the nucleus. This is consistent with previous observations that nuclear speckles, for instance, are low-density condensates with total macromolecular concentrations not dramatically different from the surrounding nucleoplasm [62, 63], and argues strongly that we are tracking a single RNA species whose hydrodynamic properties are invariant: what changes is exclusively the architecture of transient interactions it encounters. Together, these observations are consistent with a model in which NEAT1-1 outside speckles potentially alternates between nanocondensate-trapping zones and chromatin-meshwork zones as it diffuses through the nucleoplasm. Resolving this heterogeneous diffusion behavior within a single measurement is enabled by our SPAD array–based photon-resolved microscopy platform, which performs svFCS simultaneously across multiple observation volumes, thereby avoiding temporal averaging artifacts.

The lower confinement strength outside speckles compared to within them is consistent with the smaller size and reduced internal complexity of nanocondensates relative to micron-scale MLOs [64]. Nanoscale RNA-containing condensates are increasingly recognized as widespread features of the nucleoplasm [65], and several have been directly observed in the vicinity of nuclear speckles. These include transcriptional condensates concentrating RNA and RNA-processing factors at the 50–300 nm scale [66, 67], as well as RNA-containing condensates distinct from canonical speckles in the perinuclear space [68].

The close proximity of NEAT1-1 microspeckles to nuclear speckles, together with the svFCS evidence for nanoscale confinement of NEAT1-1 in the nucleoplasm, supports the hypothesis that the nucleoplasmic slow-diffusing component represents NEAT1-1 nanocondensates intended as satellite condensates that may function as MLO precursors or independent functional assemblies. Whether the interaction network of NEAT1-1 inside paraspeckles and in the nucleoplasm is shared or entirely distinct remains an open question. While this study is limited to an overexpression system, the FLAPs–FFS strategy is generalizable and provides a template for future characterization of endogenous NEAT1-1 dynamics. Full mapping of the nanoscale structures in which NEAT1-1 participates outside speckles remains challenging in living cells, and will require approaches such as diffusion-law maps across the cell [23] or single-molecule tracking combined with condensate markers. Our study opens the way to elucidating the functions of NEAT1-1 outside speckles and to mapping the nanoscale RNA-condensate landscape in living cells.

## 4. Materials and methods

### Plasmid cloning

The NEAT1-1 and biRhoBAST (2xbiRhoBAST) sequences were PCR-amplified from pcDNA5-FRT-NEAT1 and mEGFP-2xbiRhoBAST (Addgene plasmid: 196349) plasmids, respectively. PCR oligos are reported in table T1. After digestion with FastDigest PacI (Thermo Fisher Scientific, Waltham, MA, USA), NEAT1-1 and 2xbiRhoBAST were ligated. The pAAVS1-Neo-CAG-M2rtTA-H2BGFP plasmid was used as host vector upon digestion with FastDigest SalI and KpnI (Thermo Fisher Scientific). The pAAVS1-Neo-CAG-M2rtTA-H2BGFP plasmid was a gift from Roland Friedel (Addgene plasmid: 85798). NEAT1-2xbiRhoBAST sequence was digested with FastDigest SalI and KpnI (Thermo Fisher Scientific), then cloned into pAAVS1-Neo-CAG-M2rtTA. For NEAT1-only control (fig. S5), biRhoBAST repeats were removed from pAAVS1-Neo-CAG-M2rtTA-NEAT1-2xbiRhoBAST via digestion with FastDigest PacI and SalI and the linear plasmid was self-ligated following blunt-end repair. All PCR reactions were executed using Phusion™High Fidelity DNA Polymerase (M0530, New England Biolabs, Inc., Ipswich, MA, UK) following manufacturer’s protocol. PCR and restriction enzyme reactions were purified with QIAquick PCR purification kit (QIAGEN, Aarhus, DE). All the ligation reactions were executed using T4 DNA ligase (EL0011, Thermo Fisher Scientific) following manufacturer’s protocol.

### Bacterial culture

An amount of 5 *µL* ligation reaction was transformed into chemically competent TOP10*α* strain (Thermo Fisher Scientific) following manufacturer’s protocol in LB Broth with agar (L3147, Sigma-Aldrich) plates supplemented with 100 *µg/mL* ampicillin (Sigma-Aldrich). Single bacteria colonies were selected and grown overnight in LB Broth medium (L3522, Sigma-Aldrich) supplemented with 100 *µg/mL* ampicillin (Sigma-Aldrich) at 30 ^◦^C. Plasmids were purified from the bacterial culture using QIAprep Spin Miniprep Kit (QIAGEN) and quantified using a spectrophotometer (NanoDrop One, Thermo Fisher Scientific).

### Cell culture and transfection

HEK293T and HeLa cells were cultured in complete medium (DMEM, high glucose, no phenol red (Gibco™, Waltham, USA) supplemented with 100 *U/mL* penicillin/streptomycin (Gibco™) and 10% fetal bovine serum (Thermo Fisher Scientific)) at 37 ^◦^*C* and 5% *CO*_2_. All the experiments were performed by seeding cells at 50% confluency in *µ*Slide 8-well or 18-well, glass-bottom chambers (Ibidi GmBH, GE). For immunofluorescence experiments, glass-bottom chambers were coated with Poly-D-Lysine prior to seeding. Briefly, chambers were incubated for 30 min with 50 *µg/mL* Poly-D-Lysine in distilled water and rinsed three times with distilled water. Once chambers were completely air-dried, cells were seeded as explained above and cultured at 37 ^◦^C and 5% *CO*_2_. After 24 hours from seeding, cells were transfected with FuGENE® Transfection Reagent (Promega Corporation, Madison, WI, USA). For NEAT1-biRB expression, pAAV-Neo-CAG-M2rtTA-NEAT1-2xbiRhoBAST was co-transfected with mEGFP (NEAT1:GFP ratio = 1:5). For FLFS measurements performed on the custom-built microscope, EGFP was replaced by nls-EGFP (NEAT1:GFP ratio = 1:10), which served the dual role of transfection control and nuclear marker (the custom setup is not equipped with a brightfield channel for nuclear identification). The nls-EGFP plasmid was a gift from Rob Parton (Addgene plasmid: 67652). After 5 hours from transfection, medium was removed and replaced with complete medium supplemented with 1 *µg/mL* doxycycline (Sigma-Aldrich). The day after, complete medium was remove and live cell imaging or immunofluorescence were performed as decribed below.

For 1,6-hexanediol treatment, the day after transfection cells were incubated with 2% (w/v) 1,6-hexanediol (240117, Sigma-Aldrich) in complete medium for 5 min. Cells were quickly washed with DPBS. Fixation and permeabilization followed as described in *Immunofluorescence* section.

### Live cell imaging

After 24 hours from doxycycline treatment, medium was replaced with Leibovitz L-15 medium supplemented with 200 nM SpyRho 555 dye (Spirochrome AG, Walzenhausen, CH). Cells were imaged after 1 hour incubation at 37 ^◦^*C*.

### Immunofluorescence

The day after transfection cells were washed with DPBS (Gibco) and fixed with 2% paraformaldehyde in DPBS (J61899, Thermo Fisher Scientific) for 9 min. Cells were incubated with 100 mM glycine (Sigma-Alrich) in Mg-PBS (DPBS (Gibco™) supplemented with 1 mM *MgCl*_2_ (Sigma-Alrich)) for 10 min. After 2 washes in Mg-PBS, cells were incubated in blocking buffer (5% bovin serum albumin (Sigma-Alrich) in Mg-PBS supplemented with 0.1% Triton X-100 (Sigma-Alrich)) for 1 hour. Cells were incubated with 1:500 anti-NONO or anti-SC35 antibodies (Sigma-Aldrich) in blocking buffer for 1 hour. Following 3 washes in blocking buffer, cells were incubate with 1:1000 anti-mouse or anti-rabbit IgG Alexa488 or IgG Alexa647 (Thermo Fisher Scientific) in blocking buffer for an additional hour. Cells were washed with blocking buffer 3 times (5 min each) and with Mg-PBS once. Finally, cells were incubated with 0.5 *µg/mL* Hoechst-33342 (Thermo Fisher Scientific) in Mg-PBS for 10 min. Cells were washed with Mg-PBS, incubated with 100 nM SpyRho 555 dye (Spirochrome AG, Walzenhausen, CH) in Mg-PBS and imaged directly.

### Confocal imaging

Confocal imaging of live and fixed cells was performed with a spinning disk system using a Yokogawa W1 scanhead attached to a Ti2 Nikon inverted microscope. The microscope is equipped with a Zyla 4.2P camera (CMOS camera). Measurements were acquired with a Nikon Plan Apo *λ*D 100× oil objective (NA = 1.45). Imaging related to 1,6-hexanediol treatment (fig. S3) was performed using NSPARC Nikon microscope equipped with a Nikon 100× APO objective (NA = 1.25). All the images related to FLFS measurements were acquired with a custom photon-resolved laser-scanning microscope equipped with a 5×5 SPAD array detector. The microscope was introduced in the [28–30, 69] and briefly described in *Fluorescence fluctuation spectroscopy* section. Images recorded within the svFCS measurements were obatined summing all 5×5 pixel of the array detector, while images related to FLFS measurements were obtained summing the inner 3×3 pixel of the array detector.

### Image analysis

Speckle area in fig.1 was quantified via CellProfiler 4 [70]. Briefly, nuclei were segmented from DNA staining images (Hoechst-33342, 380 nm channel) to define the regions of interest. Within these nuclear boundaries, NONO-positive paraspeckles and SC35-positive microspeckles were segmented in the 488 nm channel. Independent thresholding parameters were optimized for each speckle class to take into account differences in size and signal intensity. To obtain absolute measurements, speckle areas were converted from pixels to *µm*^2^ based on the image pixel size. The corresponding CellProfiler pipelines have been deposited in the Zenodo repository (see *Data availability* section).

### Fluorescence fluctuation spectroscopy

All the fluorescence fluctuation spectroscopy experiments have been performed on a custom laser-scanning microscope designed for photon-resolved microscopy with a SPAD array detector. The microscope uses a 5×5 SPAD array detector fabricated with a 0.16 µm BCD technology [71] with the sensitive elements cooled to -15^◦^C, resulting in a reduced dark noise [69]. Briefly, the microscope excites the sample with a triggerable 485 nm pulsed laser diode (LDH-D-C-485, PicoQuant, Berlin, Germany). We coupled the laser beam into a polarization-maintaining fiber before sending it to the microscope. The laser is controlled by a laser driver, which can be synchronized via an oscillator module (PDL 828-L “SEPIA II”, Picoquant). The oscillator provides the synchronization signal to feed into the multi-channel time-tagging module to measure the histogram of the photon-arrival times needed for the fluorescence lifetime analysis. The excitation beam is deflected by a pair of galvanometric scan mirrors and focused on the sample using a 100×/1.4 numerical aperture objective lens (Leica Microsystems, Wetzlar, Germany). The fluorescence signal is collected by the same objective lens, de-scanned by the two galvanometric mirrors, and imaged on the SPAD array detector. Measurements were acquired with an average power of 1.52 *µW* and 9.95 *µW* for 488 nm and 561 nm laser lines, respectively.

### Control and data-acquisition system for photon-resolved microscopy

The microscope is controlled with the BrightEyes microscope control suite (BrightEyes-MCS) [72], which operates through a data acquisition board based on an FPGA (NI FPGA USB-7856R, National Instruments). The BrightEyes-MCS control unit drives the galvo mirrors and the axial piezo stage. It also records the signals collected by the SPAD array detector, all synchronized with the scanning beam system. The software, built on a LabVIEW system (National Instruments), utilizes the Carma application as its backbone [73]. It provides a graphical user interface for controlling microscope acquisition parameters, registering digital signals from the SPADs, and visualizing recorded signals in real-time. When fluorescence lifetime measurements are combined with imaging or correlation acquisitions, we integrate the BrightEyes-TTM module between the SPAD array detector and the FPGA-based control unit, as described in [29]. In this setup, the BrightEyes-MCS control unit supplies the synchronization signals for the scanning process (i.e., pixel clock, line clock, and frame clock) to the BrightEyes-TTM. The BrightEyes-TTM also receives synchronization signals from the pulsed laser to measure photon arrival times relative to the fluorophore excitation event.

### Experimental protocol

The detection volumes associated with the channels of the SPAD array detector were calibrated using circular scanning FCS [69]. Specifically, we performed FCS measurements of a sample of red carboxylate FluoSpheres™(F8786, 2% solids, 20 nm diameter, exc./em. 580/605 nm, Invitrogen) diluted in ultrapure water 1:5000 and drop-casted on top of a glass coverslip. The fluorescence intensity was acquired for about 120 seconds and analyzed offline. All measurements were performed at room temperature. For 561 nm channel, the detection volumes *ω*_0_ are 401, 423, and 436 nm for the SPAD array detector (respectively, from the smallest to the biggest *ω*_0_, fig. S2b).

### Fluorescence correlation calculation and analysis

We calculated the correlations directly on the lists of absolute photon arrival times [74]. For spot-variation FCS analysis, the photon lists of all relevant SPAD channels were merged, and the correlations were calculated for each observation volume. The temporal data was split into chunks of 5 or 10 s, and for each chunk, the correlations were calculated. The individual correlation curves were visually inspected, and all curves without artifacts were averaged. All the measurements were analyzed with the BrightEyes-FFS software [75]. A detailed description of the fitting functions can be found in the supporting information file and in the BrightEyes-FFS software documentation (see *Data availability*). For both single-point and circular scanning [76] FCS, the correlation curves were fitted with a 1-component or 2-component model, depending on the sample, assuming a Gaussian detection volume, as described in [69]. In the case of calibration measurements with fluorescent beads, one diffusing component model is used; in the case of SpyRho measurements, a one or two diffusing-component model was necessary (according to what stated in the manuscript). While a free-diffusion model (*α* = 1) was sufficient for the majority of datasets, an anomalous diffusion model was required for a minority of measurements where nuclear crowding hindered Brownian motion. The goodness of the fit was always checked by evaluating the reduced chi-squared and the Bayesian information criterion. For all FCS measurements acquired with the SPAD array detector, the correlation curves shown in the manuscript are related to the sum of the 25 SPAD channels. For the circular FCS measurements, the periodicity and radius of the scan movement were kept fixed while the autocorrelation amplitude, diffusion time, and focal spot size were fitted. This procedure was used for the fluorescent beads and allowed the different focal spot sizes to be calibrated (fig. S2c). For the conventional FCS measurements, the focal spot size was fixed at the values found with circular scanning FCS, and the amplitude and diffusion times were fitted. The diffusion coefficient *D_app_*can be calculated from the diffusion time *τ_D_* and the focal spot size *ω*_0_ via *D_app_* = *ω*^2^/(4*τ_D_*). For the fluorescence lifetime measurements, we calibrated the microscope by measuring the instrument response function of the complete setup (microscope, detector, and BrightEyes-TTM) using a solution of Abberior STAR 580 (Abberior GmbH, Göttingen, GE), exhibiting a fluorescence lifetime in H_2_O equal to 3.5 ns. The FLFS measurements were analyzed with a custom Python-based analysis presented in [29]. Briefly, time decay histograms of the 25 elements of the SPAD array detector were fitted individually with a mono-exponential decay function to retrieve the average fluorescence lifetime *τ_L_*. For lifetime-filtered analysis, two filter functions were computed for each detector element, corresponding to the fast- and slow-decaying components. The correlation curve for each component was then obtained by assigning a weight to every detected photon using the appropriate filter function and calculating the corresponding photon-weighted correlation, as described in [29].

### Statistical analysis

Statistical analysis was performed using GraphPad Prism, version 8.4.2 (GraphPad Software, Boston, USA). For each dataset, we performed a Shapiro-Wilk test to check the normality of the data distribution. When the hypothesis of normality was rejected, a non-parametric test (Mann-Whitney or Kruskal-Wallis tests) followed by a post hoc test (Dunn’s or Conover’s tests) was performed. When the hypothesis of normality was accepted, a T-Student test followed by a Welch’s correction was performed. All the values in the manuscript are reported as *mean* ± *SEM* .

### Data availability

The source code of BrightEyes-FFS software is available on GitHub (http://dx.doi.org/10.64898/2026.04.08.717207). All the codes used for analyzing the photon time-tagging measurements (fluorescence lifetime fluctuation spectroscopy) have been deposited in the GitHub BrightEyes-TTM repository as a part of a larger open-hardware/software project about single-photon laser-scanning microscopy based on SPAD array detector [29], and they are also available on Zenodo (https://doi.org/10.5281/zenodo.7064910). All the data, images, file measurements and analysis used for the manuscript have been deposited on Zenodo (*link available upon publication*).

## Acknowledgments

We thank all members of the Molecular Microscopy and Spectroscopy group at the Istituto Italiano di Tecnologia (IIT); Dhrisya Sathyan from the Sunbul group at the Heidelberg University for support with plasmid amplification; Dr. Dafne Campigli di Giammartino for valuable scientific input and support with grant writing; and all members of the RNA Initiative at the Istituto Italiano di Tecnologia. We also thank the Nikon Imaging centers at BioQuant (Heidelberg, Germany) and IIT (Genova, Italy), particularly Dr. Marco Scotto for support with the experiments.

## Contributions

S.Z., G.V. and M.S. conceived the idea. E.P., E.S., and G.V. developed the methodologies. G.V., E.P. and M.S. supervised and coordinated the project. S.Z. and M.O. prepared the sample. S.Z. and E.S. performed the experiments. E.S. and G.V. designed and built, with the help of E.P., the custom laser-scanning microscope. E.S. designed and implemented both the microscope control and analysis software. S.Z. analyzed the data. E.S. and E.P. helped with the data interpretation. All authors discussed the results. S.Z. wrote the manuscript with input from all authors.

