## Supplementary Information for "Mapping the diffusional landscape of short NEAT1 in living cells"

*Fluorescence correlation spectroscopy theory*

Fluorescence Correlation Spectroscopy (FCS) is a microscopy method based on the single-molecule detection of fluorescence intensity fluctuations which can be used to describe the ensemble mobility of a fluorescently tagged sample. FCS is part of a larger group of methods called fluorescence fluctuation spectroscopy. All these methods evaluate the fluorescence fluctuations to retrieve information about the mobility and concentration of the sample. In FCS, the fluorescence intensity, which is acquired in a single point of the sample space over time, is autocorrelated in time at different time delays, the so-called lag times $\tau$, and the calculated autocorrelation function, $G(\tau)$, is fitted with an analytical model describing the type of mobility of the sample inside a confined optical detection volume. For a detailed mathematical derivation of the analytical form of FCS, see Elson E. *et al.*, *Biopolymers* (1974).

For the measurements of SpyRho, we fitted the autocorrelation curves with either a model for a single component freely diffusing in a 3D environment or a model for two components freely diffusing in 3D, with a confocal observation volume. For only one diffusing component in 3D:

$$G(\tau) = \frac{1}{\langle C \rangle V_{obs}} \frac{1}{1+\frac{\tau}{\tau_D}} \frac{1}{\sqrt{1+\frac{\tau}{\tau_D}(\omega_0/z_0)^2}}, \tag{1}$$

where $\langle C \rangle$ is the average concentration of tagged molecules in the observation volume $V_{obs}$, $\tau_D$ is the average diffusion time (defined as the time point at which the autocorrelation function reaches half of its maximum amplitude), and $\omega_0$ and $z_0$ are parameters describing the observation volume (the waist and the axial profile respectively).

In confocal microscopy, $V_{obs}$ is described by:

$$V_{obs} = (\pi/2)^{3/2}\omega_0^2 z_0, \tag{2}$$

and needs to be calibrated before each measurement if the diffusion coefficient $D$ is the aim of the measurement. The diffusion coefficient can then be calculated from $\tau_D = \omega_0^2/4D$, and related to the size of the tagged molecule. If the tagged molecule is spherical, the hydrodynamic radius $r_H$ can be estimated from the Stokes-Einstein equation:

$$D = \frac{K_B T}{6\pi\eta r_H}, \tag{3}$$

with $K_B$ the Boltzmann constant, $T$ the temperature and $\eta$ the dynamic viscosity. The average number of molecules in the effective volume can be calculated from the average concentration as:

$$N = \langle C \rangle V_{eff}. \tag{4}$$

In the case of multiple diffusing components, the autocorrelation function has multiple diffusing times. Assuming that the diffusing components have the same brightness, a model for two components diffusing in 3D in a confocal volume is described by:

$$G(\tau) = \frac{1}{N}(N_1 G_1(\tau) + N_2 G_2(\tau)), \tag{5}$$

where $N$ is the average total number of tagged molecules in the observation volume, $N_1$ and $N_2$ are the percentage of number of molecules for respectively component 1 and component 2, and $G_1(\tau)$ and $G_2(\tau)$ represent the diffusion-related part of the autocorrelation function for each component. For example, for component one:

$$G_1(\tau) = \frac{1}{1 + \frac{\tau}{\tau_{D1}}} \frac{1}{\sqrt{1 + \frac{\tau}{\tau_{D1}}(\omega_0/z_0)^2}}, \tag{6}$$

with $\tau_{D1}$ the diffusing time for component one.

For the Maximum Entropy based analysis, the above concept was extended to $N_c$ components. Each fit was initialized with a uniform distribution of $N_c = 200$ diffusion-time components on a logarithmic time scale. The contribution of each component was then iteratively optimized by minimizing the sum of the squared residuals while maximizing the entropy to achieve the widest possible distribution, similarly to [77].

Calibration measurements were performed with circular scanning FCS. We acquired the fluorescence intensity over time while moving the laser beam in a circular orbit on the sample. The two parameters $\omega_0$ and $D$ that are coupled in conventional FCS through $\omega_0^2 = 4D\tau_D$ are now decoupled, since the autocorrelation function, $G(\rho, \tau)$ is a function of both spatial shifts $\rho$ and temporal lags $\tau$:

$$G(\rho, \tau) = \frac{1}{N\left(1 + \frac{\tau}{\tau_D}\right)\sqrt{1 + \frac{\tau}{\tau_D}}} \exp\left(-\frac{\rho^2}{\omega_0^2\left(1 + \frac{\tau}{\tau_D}\right)}\right). \tag{7}$$

In the case of a laser beam scanning in circles, $\rho(\tau)$ is written as:

$$\rho(\tau) = \sqrt{2R^2\left(1 - \cos\left(\frac{2\pi}{T}\tau\right)\right)}, \tag{8}$$

where $R$ is here the radius of the scan pattern, $\gamma$ is the angle over which the laser beam has traveled in a time interval $\tau$ and $T$ the time needed to scan a full circle. The parameter $\rho(\tau)$ corresponds to the distance between the two points under the angle $\gamma$. Combining Eq. 7 and 8 we obtain:

$$G(\rho, \tau) = \frac{1}{N\left(1 + \frac{4D\tau}{\omega_0^2}\right)\sqrt{1 + \frac{4D\tau}{z_0^2}}} \exp\left(-\frac{4R^2\sin^2\left(\frac{\pi}{T}\tau\right)}{\omega_0^2 + 4D\tau}\right). \tag{9}$$

Since $D$ and $\omega_0$ now appear decoupled in the equation, they can be resolved together in a single experiment.

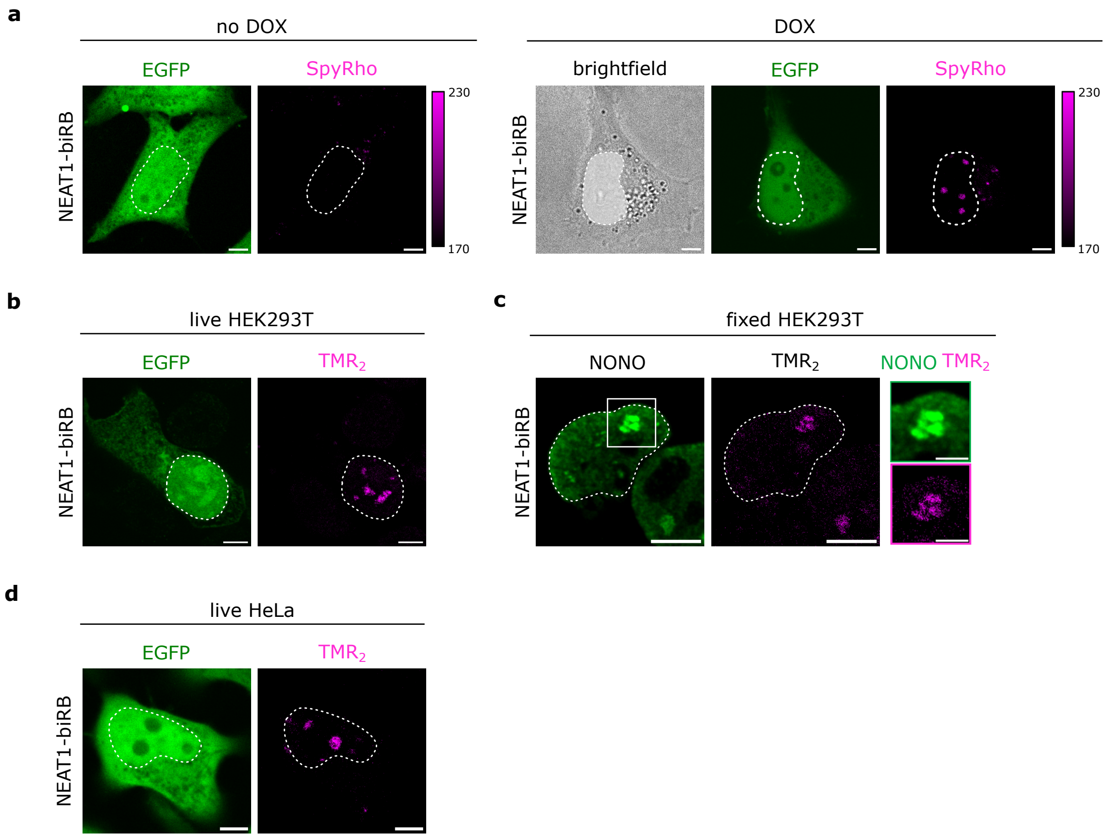


**Figure S1. Validation of NEAT1-biRB system via imaging. (a)** Brightfield and fluorescence images of transfected cells stained with SpyRho with (right) and without (left) doxycycline (DOX) induction. EGFP is used as transfection control. Figure on the right refers to 1b. **(b,d)** Representative live-cell fluorescence images of HEK293T (b) and HeLa (d) cells expressing NEAT1-biRB and stained with $TMR_2$. EGFP serves as transfection control. **(c)** Representative fluorescence images of fixed HEK293T cells expressing NEAT1-biRB. Cells were stained with $TMR_2$ (biRhoBAST) and anti-NONO antibodies (paraspeckle marker). Magnified insets (right) highlight the spatial overlap between $TMR_2$ and NONO signals. White circles indicate the nuclear boundaries. Scale bars: 5 $\mu m$.

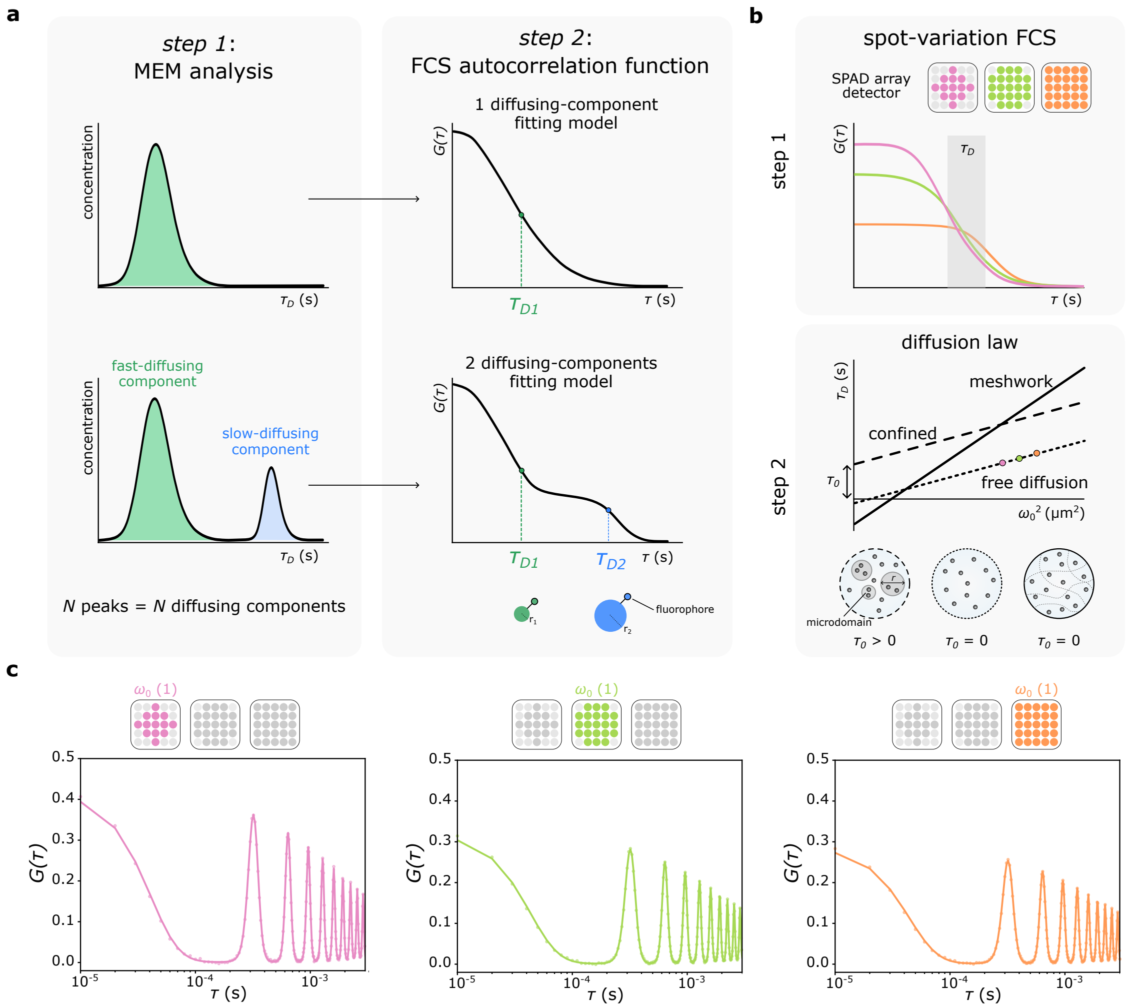


**Figure S2. Analysis workflow of FCS measurements. (a)** Sketch of the FCS analysis worflow. In step 1, the number of diffusing species is determined by the number of peaks in the $\tau_D$ distribution from the MEM analysis. In step 2, a 1 or 2 diffusing-component model is applied to $G(\tau)$ to retrieve the diffusion time $\tau_D$. **(b)** Sketch of the svFCS workflow. In step 1, FCS autocorrelation curves are retrieved from intensity traces acquired with a 5x5 SPAD array detector. By selecting different sets of single-photon detectors, 3 different observation volumes ($\omega_0$) are mimicked, resulting in 3 different curves. In step 2, diffusion times ($\tau_D$) from each ACF is used to retrieve a linear function or diffusion law. The intercept of the function ($\tau_D(0)$) determines the diffusion mode (confined, meshwork or free diffusion). **(c)** Determination of the lateral size of focal spots used for spot-variation FCS with circular scanning FCS. The panel shows a representative circular scanning FCS measurement and respective correlation curves of 20 *nm* fluorescent beads, obtained by selecting $\omega_0$ and diffusion time $\tau_\text{D}$ as fit parameters.

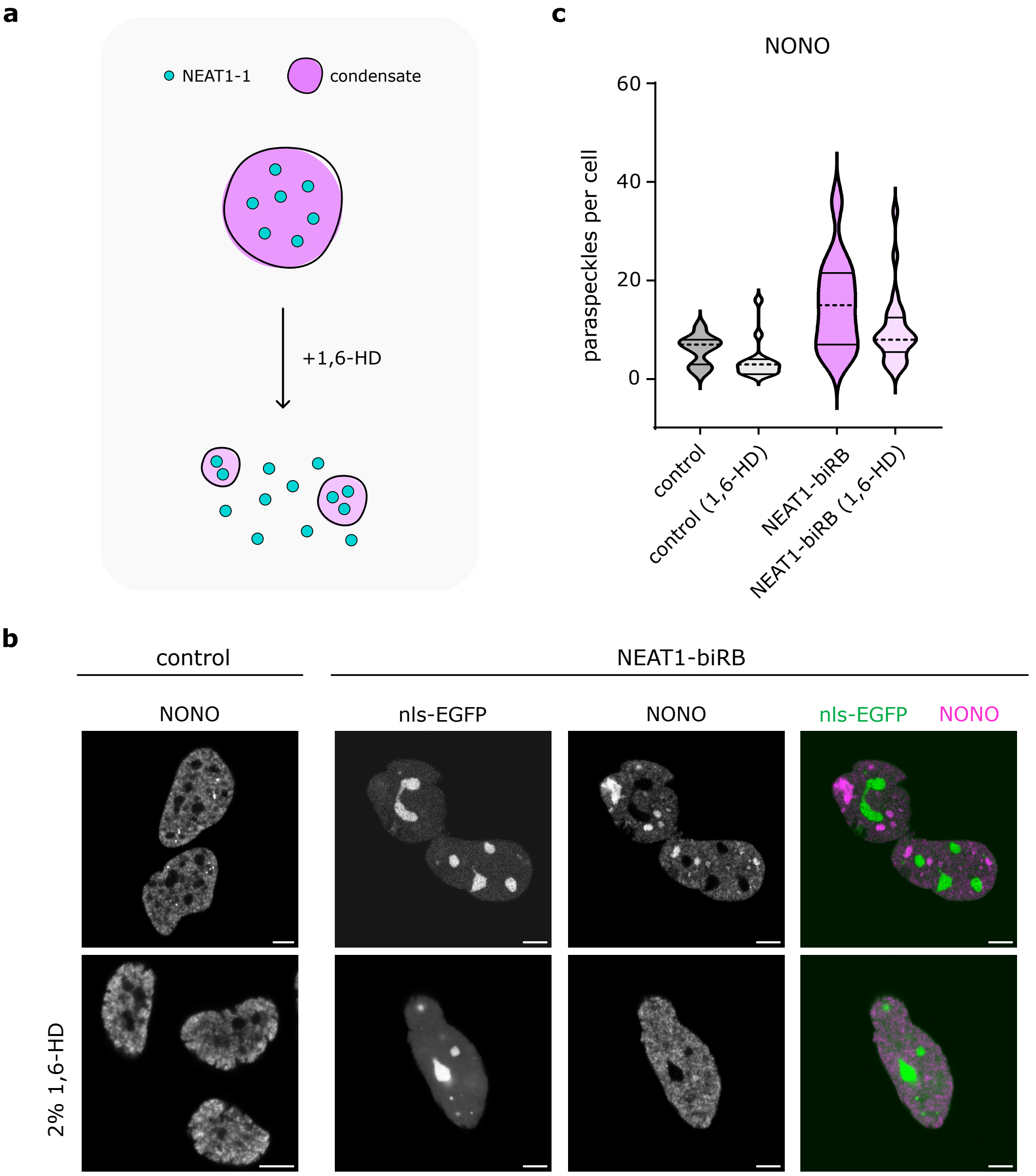


**Figure S3. NEAT1-biRB speckles retain liquid-like properties. (a)** Sketch of the effect of 1,6-hexanediol (1,6-HD) treatment, which is responsible for disruption of liquid-like condensates. **(b)** Quantification of paraspeckle (NONO) number and area in HeLa cells (control and NEAT1-biRB) before and after 1,6-HD treatment. **(c)** Representative fluorescence images of NONO immunostaining in fixed HeLa cells, both for untrasfected and NEAT1-biRB-trasfected cells. The nls-EGFP was used as transfection control. Cells are shown before (top) and after the treatment with 2% 1,6-HD (bottom). Scale bars: 5 $\mu m$.

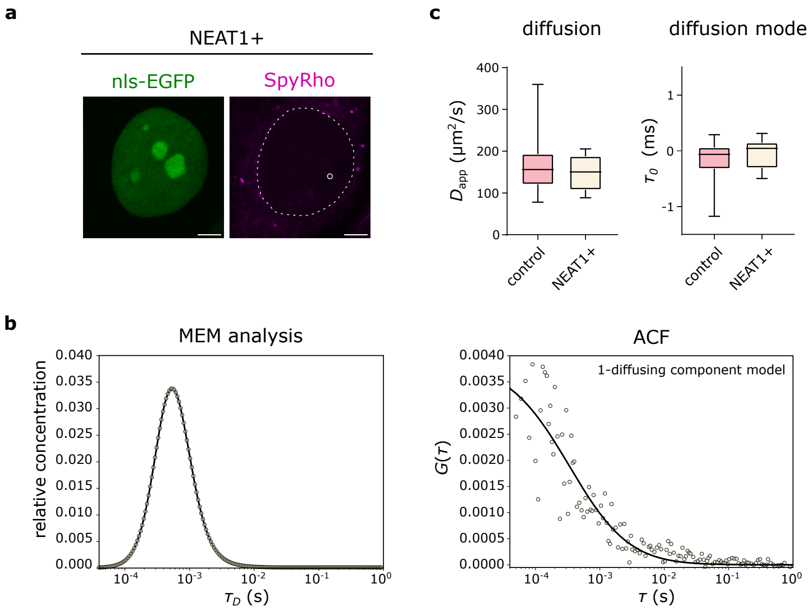


**Figure S4. Overexpression of NEAT1-1 in absence of biRhoBAST.** (a) Fluorescence images of live HeLa cells overexpressing NEAT1-1 (NEAT1+) labeled with SpyRho. Nls-EGFP serves as transfection control. A dashed circle indicate nuclear boundaries; a solid circle indicate the spot of the FCS measurement shown in (b) and (c). (b) Representative FCS measurement output within the nucleus of a NEAT1+ cell. Both MEM analysis and ACF indicate the presence of a single diffusing component. (c) Box plots of $D_{app}$ and $\tau_0$) for control and NEAT1+ cells. Both parameters are comparable across conditions, confirming that NEAT1-1 overexpression does not induce abnormal labeling or altered diffusion of SpyRho.

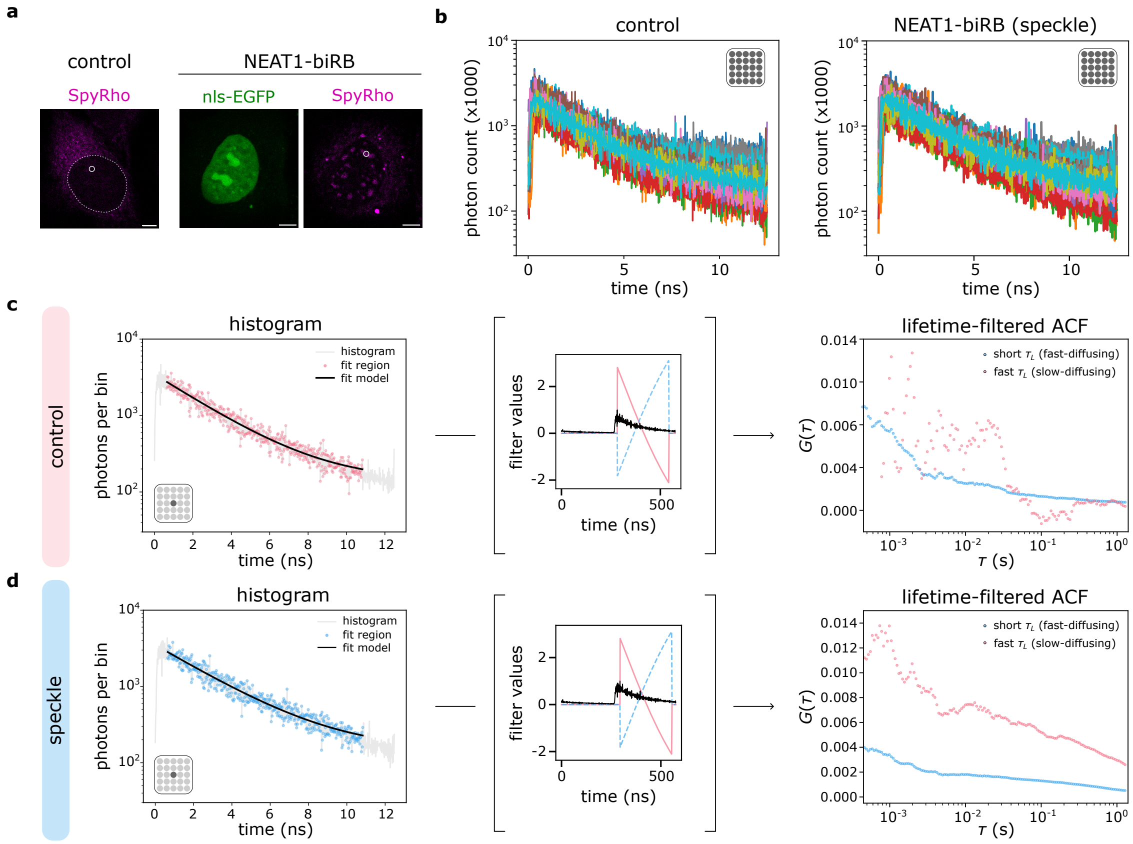


**Figure S5. Analysis workflow of FLFS measurements. (a)** Representative fluorescence images of SpyRho staining in control and transfected cells. For transfected cells, nls-EGFP serves as a transfection control. Circles indicate the specific spots (nucleoplasm in control; speckle in transfected cells) where the FLFS measurements shown in (b,c,d) were performed. Brightness and contrast were adjusted uniformly across the 561 channel for all images. Scale bar: 5 $\mu m$. **(b)** Time decay histograms for a representative FLFS measurement of SpyRho in control (left) and NEAT1-biRB-expressing (right) cells. The 25 (5x5) overlaid histograms correspond to the individual detectors of the SPAD array detector. **(c,d)** Lifetime-filtered analysis of FLFS data. For simplicity, only the histogram of the central SPAD channel is shown. Histograms are shown after dark-count subtraction, with the highlighted fitting regions in pink (control) and blue (speckle). A solid line indicates the monoexponential fitting model ($\tau_L$ = 2.67 ns for control, $\tau_L$ = 2.89 ns for speckle). The corresponding ACF after applying lifetime filters are shown on the right.

| **Primer name** | **Sequence** |
|---|---|
| NEAT1-1 (forward) | CGGGTCTAGAGGTATGTGGGAG |
| NEAT1-1 (reverse) | TCCGGGCTCCGCTAGAGCGG |
| 2xbiRhoBAST (forward) | TCCCCTTTTAATTAACGGGTCTAGAGGTATGTGGGAG |
| 2xbiRhoBAST (reverse) | GATATCGTCGACAGTCCGCTCTAGCGGAGCCCGGA |

**Table T1. PCR primers for cloning.** Sequences (5’→3’) of DNA oligos used as PCR primers for cloning of NEAT1-1 and 2xbiRhoBAST into pAAVS1-Neo-CAG-M2rtTA vector.
